# Improved Protein Encapsulation and Delivery by Lipid Nanoparticles with Refined Ionizable Lipid Content

**DOI:** 10.64898/2026.03.10.710763

**Authors:** Eimina Dirvelyte-Valauske, Modestas Mazerimas, Bozena Pavliukeviciene, Neringa Daugelaviciene, Simonas Kutanovas, Ching-Yang Kao, You-Tzung Chen, Urte Neniskyte, Rima Budvytyte

**Author notes:** Equal contributions (Eimina Dirvelyte-Valauske, Modestas Mazerimas).

## Abstract

Efficient intracellular delivery of nucleic acids, proteins, and other biomolecules is critical to advancing therapeutic strategies and genome-editing technologies. Lipid nanoparticles (LNPs) have emerged as highly promising delivery vehicles owing to their self-assembly properties, biocompatibility, and capacity to encapsulate large molecular cargos. Their biological performance is determined largely by lipid composition, which influences particle stability, cellular uptake, membrane fusion, and intracellular trafficking. In this study, we designed and optimized LNP formulations inspired by the lipid architecture of enveloped viruses. Four distinct formulations were generated and systematically evaluated in mammalian cell culture, leading to the identification of two lead candidates with superior delivery characteristics. The biodistribution and translocation properties of these formulations were subsequently assessed using an *in vitro* brain endothelial barrier model to mimic brain environment. Furthermore, we demonstrated that the selected LNPs enable efficient and functional delivery of CRISPR–Cas ribonucleoprotein complexes to mammalian cells. Together, these findings underscore the potential of rationally engineered LNPs as versatile, safe, and effective non-viral delivery platforms for advanced genome-editing applications.

## Background

The delivery of therapeutic material to target cells is a critical step for all clinical applications. DNA, mRNA, proteins, or other biomolecules are not protected from degradation and cannot cross the cell membranes on their own without active cellular uptake or physical stimulus, such as electroporation, microinjection, sonoporation, and others. However, physical transfection can harm the cells (1) or even cause cell death (2). It is also more challenging to apply these methods to 3D experiments efficiently (3). Therefore, viral vector applications are explored for efficient delivery to more complex models (4). Yet, despite their efficacy, viral vectors pose risks, including immunogenicity (5) and genotoxicity (6). Even though the leading viral delivery platform, adeno-associated viral (AAV) vectors, are widely used in preclinical studies and are even approved as therapeutics (7), their major drawback is a limited packaging capacity due to the small size of the AAV virion (8).

With the rapid expansion of CRISPR-Cas genome-editing technologies (9), there is an increasing demand for safe, versatile, and easily adaptable delivery platforms that are not constrained by cargo size and can accommodate large gene editors and their guide RNAs. Although liposomal delivery has been used in therapeutics for decades (10), lipid nanoparticles (LNPs) have recently attracted increased interest as delivery vectors due to the growing need to apply the expanding gene-editing toolbox in mammalian cells (11).

As a delivery system, LNPs offer several advantages, including self-assembly, biocompatibility, high bioavailability, ability to carry large payloads, and a range of physicochemical properties that can be controlled to modulate their biological characteristics (12). However, LNPs can be formulated with different compositions and lipid ratios, which determine their size, physicochemical properties, encapsulation efficiency, and transfection efficacy (12,13). It is known that altering the structure and composition of LNPs enables the generation of functionally distinct lipid nanoparticles for gene therapy (13). A series of selective organ-targeting (SORT) LNPs has recently been designed for tissue-specific delivery of mRNA and the CRISPR-Cas9 genome editor (14). Recently, the surface engineering of LNPs with integrated membrane proteins derived from human pluripotent stem cell-derived neurons has enabled reproducible production of efficient neuron-targeting LNPs (15). Furthermore, significant efforts have been made in the last couple of decades to understand the mechanistic role of lipid composition on membrane fusion (16). Such knowledge has guided the development of approaches to enhance intracellular delivery. One of the strategies for improving RNP delivery is enhancing cell entry by modifying LNPs with cell-penetrating peptides or targeting ligands, such as aptamers or hyaluronan (17)

Due to their easily modifiable nature, LNPs hold great potential for further development. LNP lipid formulation analysis holds significant potential to enhance its application in mammalian cell culture, 3D culture models, and whole tissues. Therefore, we aimed to develop new LNP compositions and test those already used in the clinic. In this study, we developed LNPs with improved cellular uptake by drawing inspiration from the lipid composition of enveloped viruses. These nanoparticles incorporated synthetic ionizable lipids to facilitate DNA complexation and promote endosomal escape. Additionally, a combination of natural and synthetic helper lipids was included to ensure structural integrity, provide steric stabilization, and enhance cellular uptake, membrane fusion, and intracellular trafficking. Cholesterol and PEGylated lipids were added to improve the stability, fusion, and circulation of LNPs. We assessed four different LNP compositions in a tissue culture model and selected two formulation LNPs for further investigations. To address the biodistribution of potential LNPs in the nervous tissue environment, we employed an *in vitro* brain endothelial cell-based barrier model. We also applied our proprietary LNPs for the delivery of CRISPR-Cas ribonucleoprotein complex (RNP) to mammalian cells, thereby demonstrating their potential for effective, stable, and functional delivery of gene-editing tools.

## Materials and methods

### Materials

All phospholipids: 1,2-dioleoyl-*sn*-glycero-3-phosphocholine (DOPC), 1,2-Dioleoyl-*sn*-glycero-3-phosphoethanolamine (DOPE), 1,2-dipalmitoyl-sn-glycero-3-phosphocholine (DPPC) and, 1,2-dimyristoyl-rac-glycero-3 -[maleimide (polyethylene glycol)-2000] (DMG-PEG_2000_), 1,2-Dioleoyl-3-trimethylammoniumpropane (DOTAP), cholesterol (Cho), ergosterol (Erg) and 6-((2-hexyldecanoyl)oxy)-N-(6-((-hexyldecanoyl)oxy)hexyl)-N-(4-hydroxybutyl)hexan-1-aminium (ALC-0315), were purchased from Avanti Polar Lipids (Alabaster, AL, USA). All other chemicals were used as analytical grade (Sigma-Aldrich, St. Louis, MO, USA). Ultrapure nuclease-free water (UPH_2_O) was used throughout. Alexa Fluor™ 488 conjugate of bovine serum albumin (BSA-AlexaFluor488), Invitrogen TrueGuide Cas9 protein, and sgRNAs were purchased from Thermo Fisher Scientific (Waltham, MA, USA). Cas9-GFP fusion protein was purchased from Sigma-Aldrich (St. Louis, MO, USA).

### Preparation of the LNPs

LNPs were prepared by the thin-film hydration method (18). The mixtures of 10 mM of lipids at different molar ratios were prepared in chloroform: DOPC/DOPE/Chol/DOTAP/ALC0315/DMG-PEG2000 (20/10/30/20/15/5, LNP-I), DOPC/DOPE/Erg/DOTAP/ALC0315/DMG-PEG2000 (20/10/15/15/20/15/5, LNP-II), DOTAP/Chol/DPPC/DMG-PEG2000 (46.3/42.7/9.4/1.6, LNP-III) and DOTAP/ALC-0315/DOPE/DMG-PEG2000 (24/42/30/4, LNP-IV), with LNP-IV prepared according to the previously reported formulation (19). The solvent was removed by evaporating the chloroform solution in a gentle stream of nitrogen, followed by vacuum drying for one hour. The dried film was rehydrated with the 137 mM NaCl, 2.7 mM KCl, 10 mM Na_2_HPO_4_, 1.8 mM KH_2_PO_4_ phosphate-buffered saline solution (PBS) at pH 7.4 to obtain the 1.4 mM multilamellar liposomal formulation (MLVs). The MLVs were homogenized with an ultrasonicator for 10 min, and incubated with occasional vortexing as needed until the lipid film was no longer visible. The MLVs were then extruded 21 times through a mini-extruder (Avanti Polar Lipids, Alabaster, AL, USA) equipped with a 200 nm pore-size polycarbonate membrane, resulting in a clear suspension of LNPs. LNP sizes were determined by dynamic light scattering (DLS, see below).

### Preparation of BSA-AlexaFluor488 and of RNP complex-loaded LNPs

For complex formation between lipids and BSA– AlexaFluor48 a 500 nM concentration of BSA-AlexaFluor488 was prepared in PBS at pH 7.4. The concentration ratio for complex formation between lipids and BSA–AlexaFluor488 was 4000:1. The complex between lipids and Cas9-GFP:sgRNA (RNP) complex was prepared by mixing Cas9-GFP (350 nM) and sgRNA (1050 nM) on ice for at least 20 min before use or storage at −80°C. The concentration ratio for complex formation between lipids and Cas9–GFP:sgRNA was 4000:1.

For LNPs loading, evaporated lipid films of predefined composition were resuspended in PBS at pH 7.4 containing either BSA-AlexaFluor488 or RNP complex. The mixture was sonicated for 10 min, followed by two freeze-thaw cycles. A single freeze-thaw cycle consisted of freezing the sample for 5 min at –80 °C and then thawing it for 5 min in a water bath at 37 °C. Then, 10 min sonication step was repeated, and LNPs were kept at room temperature for 10 min. The LNPs containing RNP complex or BSA-AlexaFluor488 were then extruded 21 times through a 200 nm polycarbonate membrane (Avanti Polar Lipids, Alabaster, AL, USA). Loaded LNPs with BSA-AlexaFluor488 were centrifuged at 10,000 rpm for 10 min using a 100 kDa Microcon centrifuge filter device to isolate the unloaded BSA. Loaded LNPs with RNP were centrifuged at 4000 rpm for 5 min using a 300 kDa Microcon centrifuge filter device to isolate the unloaded RNP complex. The sizes of LNPs loaded with RNP complex or BSA-AlexaFluor488 were determined by DLS (see below). LNPs were used within 24 hours or stored for 2-7 days at 4 °C to assess their stability.

### LNPs characterization by DLS

DLS was performed on a Zetasizer Nano ZS (Malvern Instruments Ltd, UK) using a laser at λ = 633 nm and a detector at scattering angle θ = 173°. The distribution of relaxation times was calculated from the field autocorrelation function g_1_(t) by applying a regularized inverse Laplace transform technique. It was then used to determine particle size distribution via the Stokes-Einstein equation.

### HPLC to define LNPs encapsulation efficacy

High-performance liquid chromatography (HPLC) analysis was performed on a PerkinElmer Flexar system consisting of a binary LC pump, a three-channel VAC degasser, an LC autosampler, an UV/VIS detector, and a fluorescence detector with a xenon lamp. UV spectrophotometric detection was carried out at λ = 280 nm. The injection volume was 50 μL. The chromatography data were acquired using Chromera software from PerkinElmer.

To determine BSA-AlexaFluor488 encapsulation efficacy, the separation was performed on a Bio SEC-3 column (4.6×150 mm, 3 μm silica absorbent particle size, Agilent) with 3 μm silica absorbent. Mobile phase was isocratic PBS at pH 7.4, with a flow rate of 0.25 ml/min and a pressure of 30.37 bar (or 440 psi). Column temperature was maintained constant at 25 °C, and excitation and emission were set at λ = 488 and λ = 520 nm, respectively. 1% (v/v) Triton X-100 was used to disrupt LNPs and release encapsulated BSA for quantification (Fig. S1A). Here, encapsulation efficacy was calculated using Eq. 1, where *C_UP+T_* denotes the BSA concentration in the purified sample, with the intensity obtained in the presence of Triton X-100 used as a baseline correction, and *C_FULL+T_* denotes the BSA concentration in the unpurified sample, with the same baseline correction as described previously.

Unlike in experiments with BSA, Triton X-100 treatment was not required to quantify RNP loading. Here, encapsulation efficacy was calculated using Eq. 2, where *C_UP_* denotes the cargo concentration in the purified LNP sample, and *C_FULL_* denotes the cargo concentration before purification. RNP concentrations were determined by HPLC using a Brownlee Bio C18 (column 4.6×150 mm, 5 μm particle size, Perkin Elmer) and an acetonitrile/water elution buffer, which itself disrupts the lipid nanoparticles (19), eliminating the need for detergent (Fig. S1B). The optimum mobile phase, consisting of acetonitrile and water (32:68, v/v), was pH-adjusted to 2.6 with 85% orthophosphoric acid. Samples were delivered via isocratic flow at 0.25 ml/min and 33.78 bar (490 psi). The column temperature was maintained at 25 °C, and excitation and emission were set at λ = 488 and λ = 509 nm, respectively.

The encapsulation efficacy was calculated as the percentage of protein incorporated into LNPs relative to the total amount of protein used, according to the following formulas:

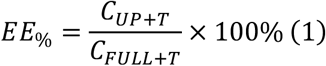

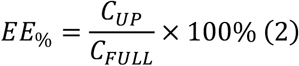

### Calcein leakage assay

To prepare calcein-loaded LNPs, the hydration buffer was replaced with 60 mM calcein solution in PBS at pH 7.4. Free calcein was separated from calcein-loaded vesicles by size-exclusion chromatography, using a chromatography column (Lenz Laborglas GmbH & Co. KG, Germany) packed with Sephadex G-50 (Sigma Aldrich, Germany).

Calcein leakage was determined using a ClarioStar Plus plate reader (BMG Labtech, Germany) in standard 96-well microplates (Corning Inc., USA) according to a previously described protocol (20). Fluorescence measurements were taken at room temperature from the bottom of the microplate every 60 minutes for 70 hours, using a 490 nm excitation filter and a 519 nm emission filter, with 8 nm and 12 nm band-pass slits, respectively. The leakage assay was performed once with four technical repeats. At the end of each measurement, the remaining calcein load was determined by adding Triton X-100 to a final concentration of 1% (v/v), which was sufficient to destroy the LNP (Fig. S1A) completely.

The data of fluorescent dye release was baseline-adjusted and normalized. Fractional permeabilization was calculated by using the equation:

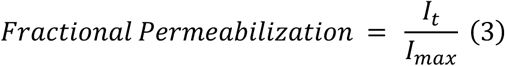

Here, *I_t_* represents fluorescence intensity at a given time point, and the *I_max_* stands for maximal fluorescence intensity after treating the sample with Triton X-100.

### Animals

Animal studies using C57BL/6J mice (RRID: IMSR_JAX:000664) were conducted in accordance with the requirements of Directive 2010/63/EU and were approved by the Lithuanian State Food and Veterinary Service (Permit No G2-92, B6-(1.9)-2653). All mice were bred and housed at the animal facility of the Life Sciences Center of Vilnius University.

### Organotypic hippocampus slice culture preparation

Organotypic hippocampal slice cultures (OHSC) were prepared using the interface method, as previously described (21,22). Briefly, 3 to 5-day-old pups of C57BL/6J mice were decapitated after cervical dislocation. The isolated brain was placed in a Petri dish filled with an ice-cold dissection medium (100 U/ml penicillin, 100 µg/ml streptomycin, 15 mM HEPES, 0.5% glucose in HBSS). The removed hippocampi were sliced at a thickness of 300 μm using a McIlwain tissue chopper. The intact slices were carefully planted on prepared 0.4 μm 30 mm diameter cell culture inserts with PTFE membrane (Merck Millipore) with maintaining medium (25% 1×BME, 25% horse serum, 5% 10×MEM, 100 U/ml penicillin, 100 µg/ml streptomycin, 2 mM GlutaMAX, 0.65% glucose, 9 mM sodium bicarbonate in ddH_2_0) and maintained in 5% CO_2_ incubator at 37 °C. The medium was changed 24 h after the plating and every 2–3 days during culture maintenance.

### OHSC treatment with LNPs

OHSCs were treated at 2 days *in vitro* (DIV2) with 350 nM BSA-AlexaFluor488 encapsulated in LNPs. A 10 μl droplet of the LNP suspension was carefully applied directly onto the surface of each slice. After 24 h, the slices were imaged using an Olympus IX83 widefield microscope with a 10×/0.3NA air objective. Fluorescent images -obtained with a FITC filter cube and phase contrast images were acquired on an Olympus IX83 microscope (Germany). Technical replicates in each experiment included three independent slices.

### Cultivation of mammalian cell lines

Human embryo kidney HEK293T/17 (RRID: CVCL_0063) and mouse brain endothelial bEnd.3 (RRID: CVCL_0170) cell lines were purchased from American Type Culture Collection (ATCC, VA, USA). Cells were maintained in T25 flasks (Corning, USA) with Dulbecco’s Modified Eagle’s Medium (DMEM) supplemented with GlutaMAX™, 10% fetal bovine serum (FBS), 100 U/ml penicillin, and 100 µg/ml streptomycin at 37 °C and 5% CO_2_. Cells were harvested using TrypLE™ for HEK293T/17 cells or 0.25% trypsin-EDTA for bEnd.3 cells. All cell reagents were purchased from Thermo Fisher Scientific (USA).

### *In vitro* endothelial cell barrier model

Mouse brain endothelial bEnd.3 cells were seeded onto a 6.5 mm Transwell® Polyester Membrane Insert with 0.4 µm pores (Corning, USA) placed into a 24-well plate at a density of 8×10^4^ cells/cm^2^. After a week, to confirm that the cells formed a tight monolayer, bEnd.3 cells were fixed with 4% PFA in PBS for 20 min. at room temperature. After rinsing with PBS, residual PFA was quenched with 30 mM glycine in PBS, and the cells were washed twice with PBS. To visualize the expression of tight junction proteins, fixed cells were permeabilized and blocked with 0.2% Triton X-100 and 3% BSA in PBS for 30 min. Afterwards, cells were incubated with primary antibodies: 1:200 polyclonal rabbit anti-ZO-1, 1:200 anti-claudin-5, and 1:50 anti-occludin (all from Thermo Fisher Scientific, USA) in PBS overnight on a shaker at 4 °C. Following three washes with PBS, the cells were incubated with secondary 1:500 goat anti-rabbit AlexaFluorPlus594 antibodies at 1:500 (Thermo Fisher Scientific, USA) and phalloidin-iFluor488 at 1:1000 (Abcam, UK) in PBS for 1 h on a shaker at room temperature. The nuclei were counterstained with DAPI at 1 μg/ml (Sigma, USA) for 10 min. at room temperature. After three more washes with PBS, cells were imaged using a 60×/1.20NA oil objective on an Olympus IX83 widefield microscope (n = 4-6 images for each well, all conditions were repeated in triplicate).

### LNPs transcytosis on an *in vitro* barrier model

Mouse brain endothelial bEnd. 3 cells were grown on Transwell® inserts, as described above, using medium without phenol red, to avoid interference with HPLC experiments. To capture transcytosis of LNPs, HEK293T/17 cells were seeded in a 24-well plate with a transwell at a density of 1.85×10^4^ cells/cm^2^. The culture medium was changed every 2-3 days. After a week, 35 nM of BSA-AlexaFluor488 encapsulated in LNPs were added to the top compartment of the transwell. Compartmentalized media were analysed by HPLC analysis after 24h, 48h, or 72h. At the same time, both bEnd.3 cells on the inserts and HEK293T/17 cells at the bottom of the wells were fixed for microscopy analysis using 4% PFA in PBS for 20 min at 37 °C. After rinsing with PBS, residual PFA was quenched with 30 mM glycine in PBS, and the cells were washed twice with PBS. All nuclei were stained with DAPI (1 μg/ml) for 10 min at room temperature and then washed 3 times. Cells were imaged using DAPI and FITC fluorescence filter cubes with 20×/0.45NA air objective on the Olympus IX83 widefield microscope (n = 4-6 images for each well, all conditions were repeated in duplicates).

### LNP delivery of CasG-GFP and sgRNA ribonucleoprotein

To apply LNPs for the delivery of ribonucleoprotein complexes (RNPs), HEK293T/17 cells were plated at a density of 2.2×10^4^ cells/cm^2^ onto 24-well plates (Corning, USA) that were pre-coated with poly-L-lysine (0.001% in PBS) for at least 20 min at room temperature. The next day, Cas9-GFP and sgRNA complexes encapsulated in LNPs were added to HEK293T/17 cells, using a Cas9 concentration of 9-35 nM. HEK293T/17 cells were incubated with RNP-loaded LNPs for 3-24 hours. Then, cells were washed with PBS to remove residual LNPs, fixed with 4% paraformaldehyde (PFA) in PBS for 20 min at room temperature. After rinsing with PBS, residual PFA was quenched with 30 mM glycine in PBS, and the cells were washed twice with PBS. Cell nuclei were stained with DAPI (1 μg/ml in PBS) for 10 min at room temperature and then washed three times with PBS. Cells were imaged under the Olympus IX83 widefield microscope with 20×/0.45NA air objective using DAPI and FITC fluorescence filter cubes (n = 4-6 images for each well, all conditions were repeated in triplicate).

### Quantification of LNP delivery efficacy

Microscopy images were analyzed using the open-source CellProfiler software (RRID: SCR_007358) (23). Cell nuclei were identified using the DAPI channel and the *IdentifyPrimaryObjects* function. The cell contour was reproduced using *IdentifySecondaryObjects* function to identify the cell cytoplasm compartment. The DAPI channel was used as a reference to expand the distance by 20 pixels. The integrated intensity of GFP was measured in each cell by the *MeasureObjects* function. Outliers were identified in GraphPad Prism (version 9.5.1 for Windows, GraphPad Software, San Diego, California, USA) and were excluded from further analysis. To account for autofluorescence in the GFP channel, the mean GFP fluorescence intensity in control images was used as a threshold to identify true GFP signal in cells. The percentage of GFP-positive cells was calculated, and data were presented as mean ± standard error of the mean (SEM) from 3 independent biological replicates, unless otherwise indicated.

### T7 endonuclease I (T7E1) assay to determine genomic cleavage

To assess genomic cleavage activity of LNP-delivered RNPs, we used TrueCut v2 Cas9 protein in an RNP complex with three different synthetic sgRNAs targeting *WTAP*, *RUNX1*, and *CDK4*. Three different RNPs were assembled and used either for LNP encapsulation or for commercial lipofection using Lipofectamine CRISPRMAX (Invitrogen). HEK293T/17 cells (2.6×10^4^ cells/cm^2^ density in 24-well plates) were transfected with 15 nM concentration of RNPs encapsulated in LNPs or with Lipofectamine CRISPRMAX following the manufacturer’s recommendations. After 48 h, cells were collected for T7E1 assay using GeneArt™ Genomic Cleavage Detection Kit (Thermo Fisher Scientific) according to the manufacturer’s recommendations. In short, cells were lysed, targeted gene fragments (approximately 500-700 bp) were amplified with PCR (primers: CDK4 5’-Gcacagacgtccatcagcc-3’, 5’-Gccggccccaaggaagactgggag-3‘; RUNX1 5‘-CCTCTTCCACTTCGACCGAC-3‘, 5‘-TGTGACCTACTATGGCTTTATGAG-3‘; WTAP 5‘-CCAGGAAGAAATTCGCCTGTTA-3‘, 5‘-TGATTACATCCTCAGACACCAA-3‘). Then, PCR fragments were denatured and reannealed to form heteroduplexes, which were digested by the T7E1 detection enzyme. DNA fragments stained with GelRed (Sigma) dye were fractionated in 1,5% agarose gel in TAE buffer, running electrophoresis for 70 min at 50 V, and imaged on ChemiDoc™ (Bio-Rad) imager. Parental and cleaved DNA bands were measured to assess T7 endonuclease I cleavage efficacy using Fiji software(24). Target lanes were selected using a rectangular tool, and lane profile plots were generated using *Mark First Lane*, *Mark Next Lane*, and *Plot Lanes* commands. The line-drawing tool was used to draw baselines to define each peak as a closed area. Peak areas were measured by clicking inside of each one with the wand tool. The equations (4) and (5) were used to calculate cleavage efficacy.

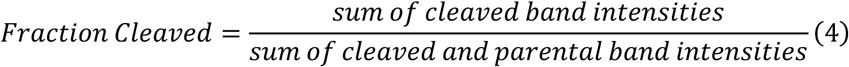

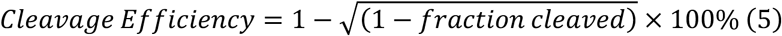

### Statistical Analysis

Statistical analysis was conducted with GraphPad Prism version 8.4.3 for Windows (GraphPad Software, USA). A significance level (α) of 0.05 was selected for the study. Data normality was assessed using the Shapiro-Wilk test. For comparing groups with normally distributed data, one-way ANOVA with the Tukey’s *post hoc* test or two-way ANOVA with *post hoc* Holm-Šídák test was employed. Statistical analysis of LNPs hydrodynamic radius measurements was done by using ordinary one-way ANOVA with Welch’s correction for unequal variances, followed by multiple comparisons of group means. To evaluate the statistical significance of hydrodynamic radius changes between BSA-AlexaFlour488 loading steps, three-way ANOVA with Tukey’s multiple comparisons was employed.

## Results

### Characterization of different formulations of LNPs

To identify the optimal content of ionizable lipids in LNPs, four formulations (LNP-I to LNP-IV) were produced (Fig. 1A). LNP-I and LNP-II were designed to test novel lipid combinations, with LNP-I unique for its reduced content of ionizable lipids, and LNP-II incorporating ergosterol as a partial cholesterol substitute. Ergosterol may alter LNP morphology by reducing lipid packing order and thereby promoting endosomal membrane fusion (25). LNP-I and LNP-II contain synthetic cationic or ionizable lipids (DOTAP and ALC-0315) to facilitate DNA complexation and promote endosomal escape. Both formulations also include a combination of natural and synthetic helper lipids (e.g., DOPC, DOPE) to maintain structural integrity, provide steric stabilization, and enhance cellular uptake, membrane fusion, and intracellular processing. Cholesterol and PEGylated lipids are incorporated to improve particle stability, promote membrane fusion, and prolong circulation (26). In contrast, LNP-III and LNP-IV were comparatively streamlined: LNP-III was dominated by DOTAP (46.3%) and cholesterol (42.7%), while LNP-IV consisted primarily of ALC-0315, cholesterol, and DOPE (Fig. 1A). LNP-III and LNP-IV were included as reference formulations based on previously reported lipid compositions and ratios. LNP-III followed previously reported molar ratios (27), but the ionizable lipid was replaced with the cationic lipid DOTAP to assess the importance of ionizable lipids for enhanced LNP delivery. LNP-IV followed the previously published 28M composition (28).

**Fig. 1.**
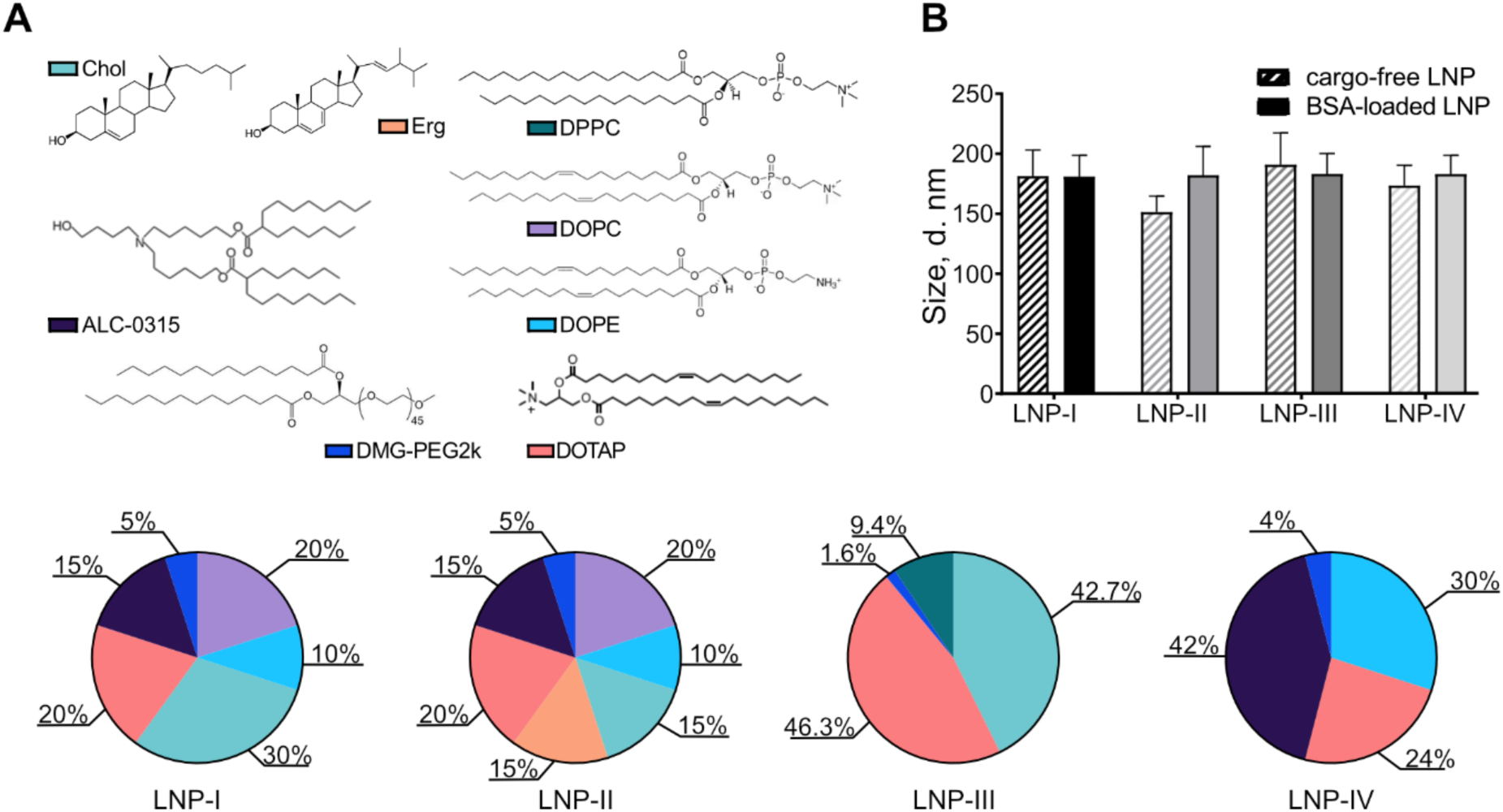
Lipid composition and size distribution of cargo-free and BSA-loaded lipid nanoparticles (LNPs). **(A)** Pie charts show the molar composition of the four LNP formulations (LNP-I to LNP-IV), with individual lipid components indicated by color codes. **(B)** Hydrodynamic diameters of cargo-free and BSA-loaded LNPs. All formulations were prepared using the thin lipid film hydration method and analyzed by dynamic light scattering (DLS, intensity-weighted). Data are shown as mean ± SEM from 13 technical replicates.

BSA was chosen as a model protein cargo due to its stability, accessibility, relatively low cost, and ease of fluorescent detection after conjugation with Alexa Fluor 488. These characteristics make BSA a practical surrogate cargo for evaluating nanoparticle formulation, processing, encapsulation, and stability under controlled conditions. The calibration curve for BSA quantification by HPLC and representative chromatograms of BSA are shown in Fig. S2 A, B. To assess whether differences in LNPs composition or BSA loading influenced particle size, the hydrodynamic diameters of the nanoparticles were measured (Fig. 1B). Across all LNPs formulations, mean diameters ranged from 151.4 to 190.8 nm under both cargo-free and BSA-loaded conditions. Importantly, the presence of BSA cargo did not significantly alter nanoparticle diameter relative to cargo-free controls, indicating that encapsulation does not grossly perturb the structural organization of LNPs at the measured scale. Taken together, these findings demonstrate that despite their distinct compositional profiles, all four formulations yield LNPs of similar sizes under both loaded and unloaded conditions, providing a consistent baseline for further analysis of processing, stability, and encapsulation.

Encapsulation efficiencies of the four formulations are summarized in Table 1. Among them, LNP-I showed the highest encapsulation efficacy of 68.2%, while LNP-II, LNP-III, and LNP-IV reached 57.2%, 52.3%, and 55.3%, respectively. Only Triton-treated BSA-loaded LNP samples showed detectable protein peaks, indicating that loaded LNPs do not leak BSA (Fig. S3 A-D).

**Table 1.** Encapsulation efficacies of BSA-AlexaFluor488 in different LNP formulations.

| Composition | Encapsulation efficacy, % |
| --- | --- |
| LNP-I | 68.17 |
| LNP-II | 57.24 |
| LNP-III | 52.29 |
| LNP-IV | 55.27 |

### LNPs’ formulation impacts mammalian cell transfection

To assess whether four different LNP formulations have different capabilities to transfect mammalian cells, we treated a 3D model of mouse organotypic hippocampal slice cultures (mOHSC) with BSA-AlexaFluor488 encapsulated in LNP-I, LNP-II, LNP-III, or LNP-IV (Fig. 2). Widefield microscopy images of live mOHSC revealed that LNP-III nanoparticles were highly adhesive and exhibited nonspecific adsorption to the brain tissue and to the insert membrane, making it unsuitable for delivery tool development. LNP-I, LNP-II, and LNP-IV did not exhibit unspecific attachment, delivering fluorescently labelled BSA exclusively to the tissue. However, LNP-IV-treated slices exhibited significantly lower fluorescence signal compared to LNP-I and LNP-II (Fig. 2). Therefore, LNP-I and LNP-II were selected for further development of delivery LNPs.

**Fig. 2.**
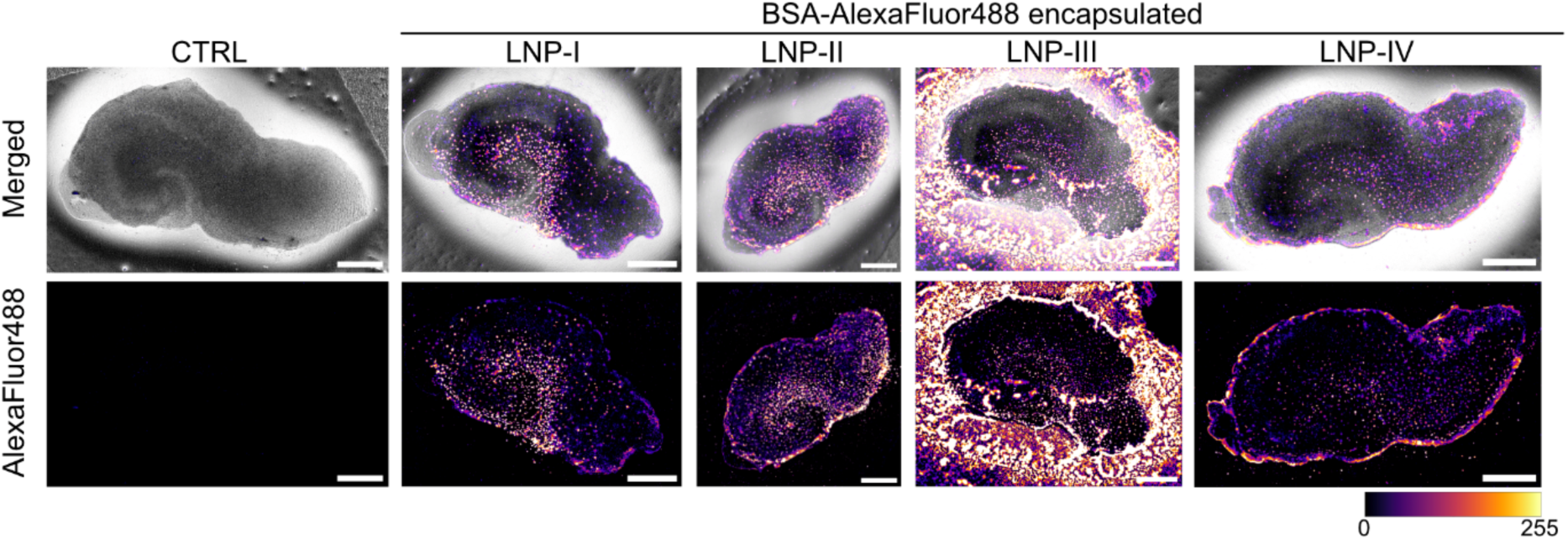
Comparison of LNP formulations for transfection in tissue culture. Widefield microscopy images of mouse organotypic hippocampal slice cultures (mOHSC) treated with bovine serum albumin (BSA) conjugated to AlexaFluor488 and encapsulated in different LNP formulations (LNP I–IV). Non-treated mOHSCs serve as a control. Scale bar 200 μm.

### Physical properties of LNP-I and LNP-II

As noted above, LNP-III and LNP-IV were primarily included as reference formulations derived from the literature and served as comparative models rather than candidates for further optimization. To evaluate how stepwise processing influenced the physical properties of LNPs, LNP-I and LNP-II were designed as novel, modifiable formulations and selected for detailed analysis at successive preparation stages (Fig. 3). In the initial resuspended (RE) and freeze– thawed (FT) states, BSA-loaded nanoparticles exhibited significantly larger diameters compared with their cargo-free counterparts (three-way ANOVA with Tukey’s multiple comparisons, *p* < 0.0001). This effect was attenuated after extrusion (EX), which reduced particle sizes and minimized the difference between loaded and cargo-free samples. Ultracentrifugation (UC) did not further reduce the mean size of LNPs compared with EX, but removed residual aggregates and debris, yielding the most homogeneous preparations, while the debris fraction (DE) consisted predominantly of particles near the detection thresholds. Statistical comparisons confirmed that LNPs’ size decreased progressively with each processing step (Fig. 3A). RE and FT samples exhibited significantly larger diameters compared with UC LNPs (*p* < 0.0001). EX resulted in a marked reduction in the mean diameter of LNPs, and although UC did not further decrease it, this step was essential for purification and debris removal. Accordingly, no significant differences were detected between UC and EX samples for most comparisons, whereas both groups differed from RE and FT (*p* < 0.0001). These results indicate that cargo loading increases initial LNPs diameters, but EX and UC largely normalize the distribution of LNPs’ sizes, with EX being the critical homogenization step and UC serving as a complementary purification step.

**Fig. 3.**
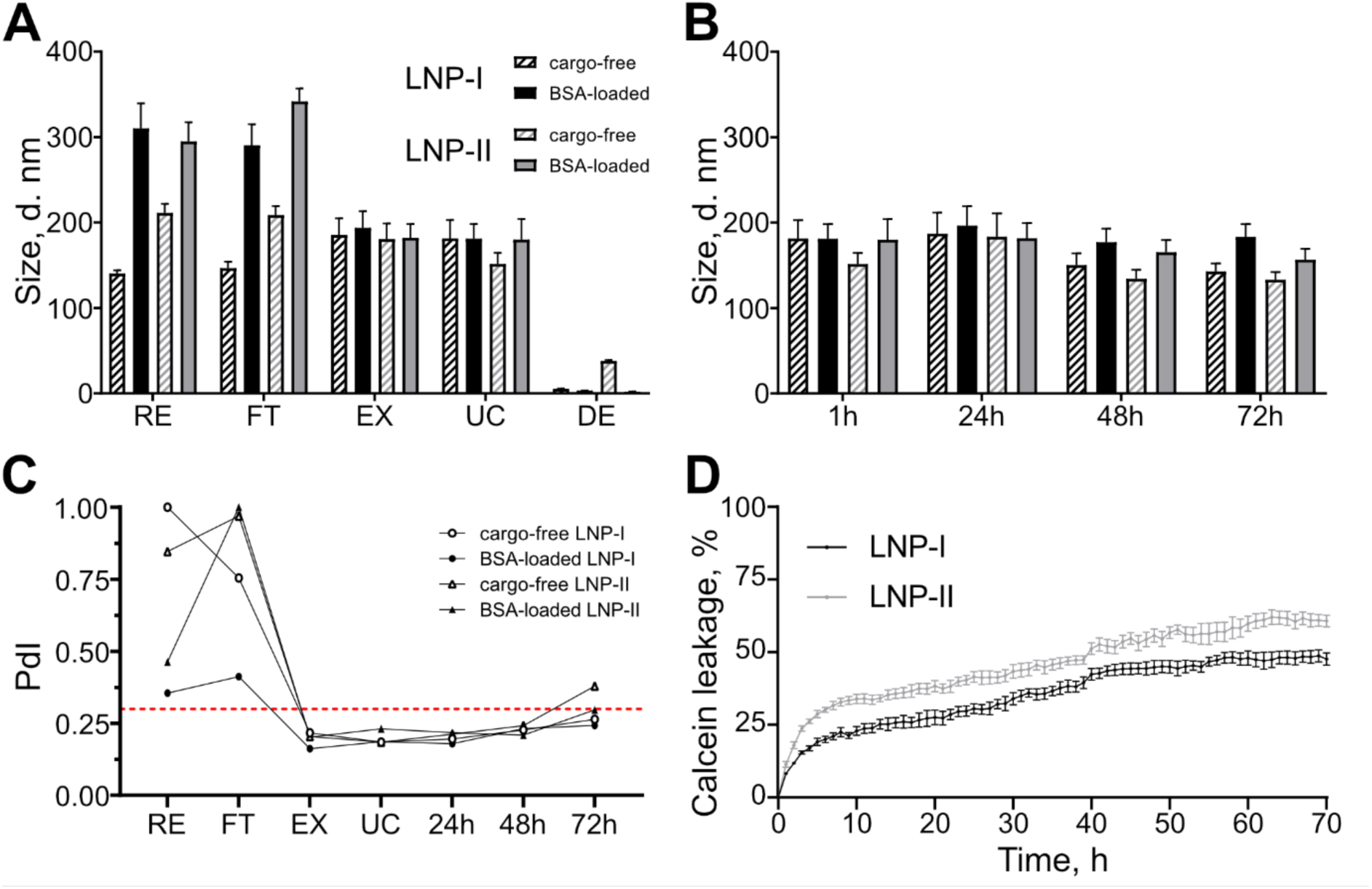
Size evolution, polydispersity, and membrane integrity of LNP-I and LNP-II formulations. **(A)** Hydrodynamic diameters of LNP-I and LNP-II, either cargo-free or BSA-loaded, measured after each preparation step, determined by dynamic light scattering (DLS, intensity-weighted). Here, RE stands for resuspended, FT – freeze-thawed, EX – extruded, UC – ultracentrifuged, and DE - debris. **(B)** Size evolution of the same formulations over time (1 h, 24 h, 48 h, and 72 h). **(C)** Polydispersity index (PdI) changes corresponding to preparation steps and time points shown in panels A and B. Data in panels A–C represent mean ± SEM of 13 technical replicates per condition, measured by dynamic light scattering (DLS). (D) Calcein leakage assay, indicating calcein leakage as a percentage of total calcein released after treating LNP formulations with 1% (v/v) Triton X-100. Data in panel D represent mean ± SEM of 4 technical replicates per condition, ClarioStar Plus plate reader.

LNP preparations were stable, retaining consistent diameters of both cargo-free and BSA-loaded LNP-I and LNP-II over 72 h storage at 4 °C, with no significant differences observed over time (three-way ANOVA with Tukey’s multiple comparisons, Fig. 3B). Polydispersity index (PDI) values were initially high in the RE and FT samples, consistent with broad particle distributions, but decreased sharply after EX and UC, reaching values below 0.25 (Fig. 3C), which is indicative of homogeneous LNPs populations (29). During subsequent storage, PDI values remained stable within this range, further supporting the conclusion that EX and UC steps are sufficient to produce stable LNPs suspensions.

Finally, the calcein leakage assay was used to evaluate cargo retention in different LNP formulations (Fig. 3D). Calcein release was consistently higher for LNP-II, reaching 62.5% after 70 hours. In contrast, LNP-I leakage did not exceed 49.0% over the same period. The higher leakiness of LNP-II is likely due to ergosterol, which is expected to reduce bilayer ordering and weaken membrane stability, thereby accelerating the release of encapsulated content.

Overall, this functional characterization revealed how lipid composition affects LNPs’ stability. In particular, LNP-II, which incorporates ergosterol, exhibited reduced durability as evidenced by increased calcein release over time compared with LNP-I. While both formulations retained comparable sizes after preparation and storage (Figs. 1B and 3A–B), LNP-II released its encapsulated content more rapidly, suggesting that ergosterol substitution compromises membrane integrity. These findings indicate that lipid composition has a critical impact on both encapsulation efficacy (Table 1) and retention stability (Fig. 3D), with LNP-I showing the most favorable performance among the tested formulations.

### LNP-I and LNP-II do not permeate the blood-brain barrier

LNPs are commonly used for various *in vivo* studies, usually to target the periphery after systemic delivery (14). Due to the lack of studies investigating LNPs’ potency to permeate the blood-brain barrier, we aimed to assess our proprietary LNPs on an *in vitro* transcytosis model (Fig. 4). Our experimental model consisted of bEnd.3 mouse brain endothelial cells that were cultivated on a porous membrane transwell insert until a tight monolayer formed (Fig. 4A). The expression of tight junctional proteins ZO-1, claudin-5, and occludin, as well as phalloidin-AlexaFluor488 labelling of F-actin in bEnd.3 cells confirmed that endothelial cells formed an intact barrier (Fig. 4B). ZO-1 and claudin-5 stained the intercellular junctions, while occludin remained mostly intracellular. To assess whether LNP-I and LNP-II cargo can be transcytosed across the barrier, LNPs loaded with BSA-AlexaFluor488 were added to the top transwell, and the LNPs in the bottom fraction were identified as transcytosed particles (Fig. 4C-F). To evaluate whether intact LNPs can cross the bEnd.3 layer, HEK293T/17 cells were seeded on the bottom of the well, acting as a target for transcytosed LNPs. After 48 hours of incubation, almost no BSA-AlexaFluor488 fluorescence was observed inside the HEK293T/17 cells, comparable to that observed in the unencapsulated BSA-AlexaFluor488 control (Fig. 4C, D). This indicated that cargo was not delivered across the barrier, suggesting that LNP-I and LNP-II were not transcytosed. Comparison between the full transcytosis model and the insert-alone condition revealed a statistically significant difference for LNP-I and a near-significant trend for LNP-II and BSA-AlexaFluor488. Importantly, both LNP formulations permeated the insert alone, with approximately 20% of GFP-positive HEK293T/17 cells, indicating that nanoparticle transport was not limited by the transwell membrane’s pore size (Fig. 4C, D).

**Fig. 4.**
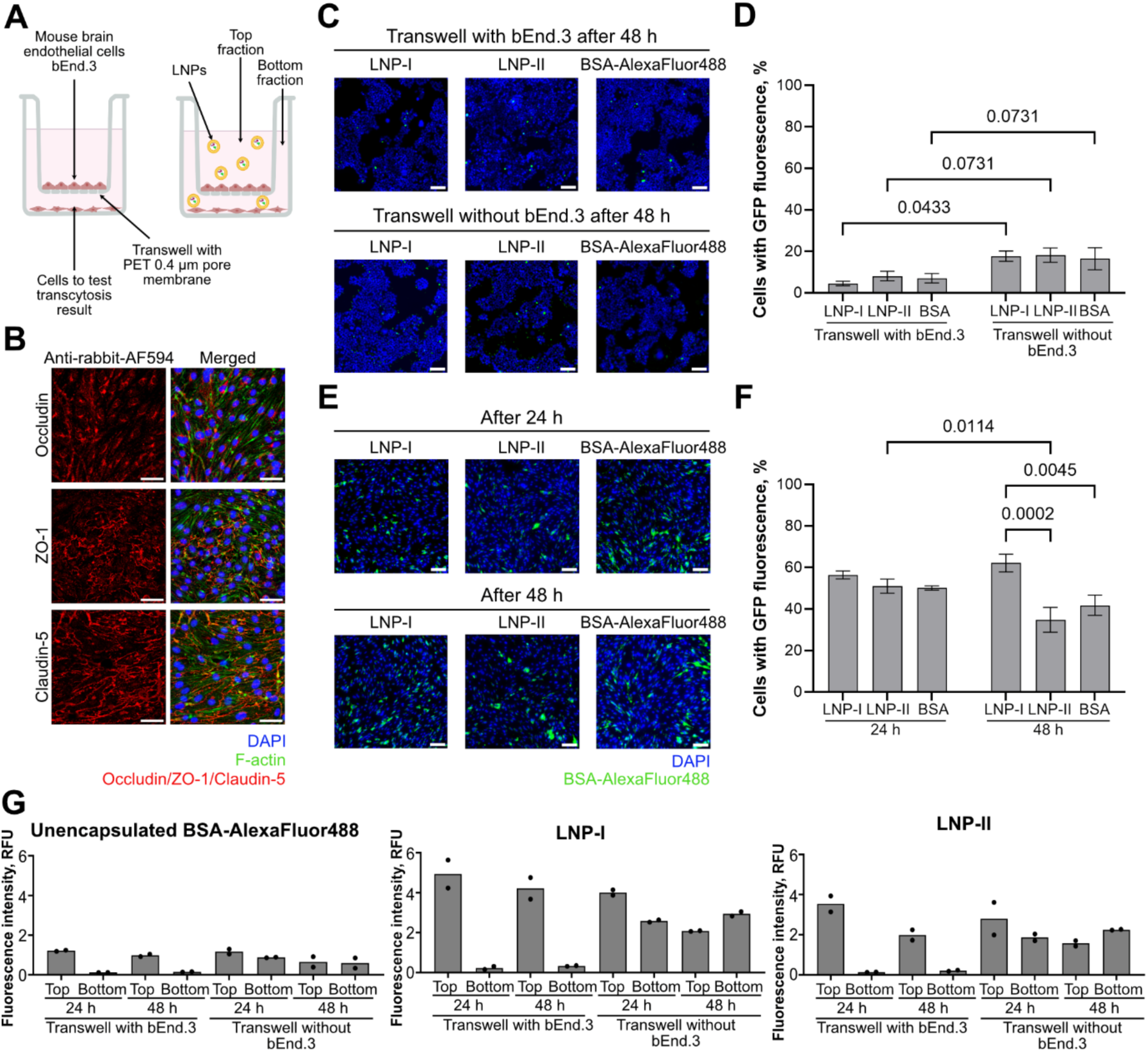
LNP transcytosis through an *in vitro* endothelial cell barrier model. **(A)** Scheme of an *in vitro* brain endothelial cells transcytosis model with bEnd.3 cell line monolayer on a porous membrane insert separating the medium into top and bottom compartments. HEK293T/17 cells are cultured in the bottom compartment to assess transcytosis across the endothelial monolayer. **(B)** Immunofluorescence images of bEnd.3 cells showing the expression of tight junction proteins occludin, ZO-1, and claudin-5. Scale bar 50 μm. **(C)** Widefield microscopy images of HEK293T/17 cells in the bottom compartment following treatment via transcytosis with LNP-I or LNP-II encapsulating BSA-AlexaFluor488. Unencapsulated BSA-AlexaFluor488 serves as a control. Scale bar 100 μm. **(D)** Quantification of GFP-positive HEK293T cells after treatment with LNPs and BSA-AlexaFluor488 control in the transcytosis model. **(E)** Widefield microscopy images of bEnd.3 cells on the inserts after treatment with LNP-I or LNP-II encapsulating BSA-AlexaFluor488. Unencapsulated BSA-AlexaFluor488 serves as a control. Scale bar 100 μm. **(F)** Quantification of GFP-positive bEnd.3 cells after LNPs and BSA-AlexaFluor488 control treatment in the transcytosis model at 24 and 48 h. **(G)** HPLC analysis of top and bottom fractions from the transcytosis model after application of either unencapsulated BSA-AlexaFluor488 or BSA-AlexaFluor488 encapsulated in LNP-I or LNP-II to the top compartment. Data are presented as mean ± standard error of the mean (SEM) from 3 independent biological replicates (D, F), or as means with individual values from two technical replicates shown as dots (G). Means were compared by two-way ANOVA and *post-hoc* Holm-Šídák test. *p*-values < 0.05 were considered significant.

Because we did not observe transcytosis, we evaluated whether LNP-I and LNP-II entered bEnd.3 cells on the transwell membrane by assessing the fluorescence inside the endothelial cells (Fig. 4E, F) and performed HPLC on the media fractions above and below the membrane with or without the bEnd.3 cells on the insert (Fig. 4G). We found that unencapsulated BSA- AlexaFluor488 control adsorbed to endothelial cells or was actively taken up by them (Fig. 4E), but was not transcytosed (Fig. 4C, D). After 24 h of LNPs and BSA-AlexaFluor488 treatment, about 50 to 60% of bEnd.3 cells were GFP-positive (Fig. 4F). In contrast, after 48 hours, bEnd.3 brain endothelial cells treated with LNP-I showed increased cargo fluorescence relative to LNP-II and unencapsulated BSA-AlexaFluor488 (Fig. 4F), suggesting enhanced transfection efficiency of LNP-I. HPLC results confirmed the widefield microscopy data: both LNP variants exhibited elevated fluorescence in the bottom fraction only when the model without endothelial cells was used (Fig. 4G). Free BSA-AlexaFluor488 was not transcytosed by bEnd.3 cells, but diffused through the insert without the cells (Fig. 4G).

### Comparison of LNPs formulations and their cargo-dependent size

The growing need for cellular delivery of state-of-the-art genome editors has recently stimulated the field of LNP development. To demonstrate the capabilities of the LNPs for efficient, stable, and functional delivery of gene-editing tools, the encapsulation and delivery performance of the LNP formulations were evaluated using a Cas9 protein complexed with sgRNA (RNP). RNP loading did not change the hydrodynamic diameters of LNP-I and LNP-II particles (Fig. 5). Consistent with earlier results (Fig. 1B), both LNP formulations maintained mean diameters in the range of ∼145–185 nm regardless of cargo type. Importantly, RNP loading did not significantly alter nanoparticle size compared with either cargo-free or BSA-loaded states, confirming that encapsulation of the larger RNP complex did not destabilize or expand the particle structures. This indicates that both LNP-I and LNP-II can accommodate Cas9 RNP complexes without significant perturbations to their nanoscale architecture.

**Fig. 5.**
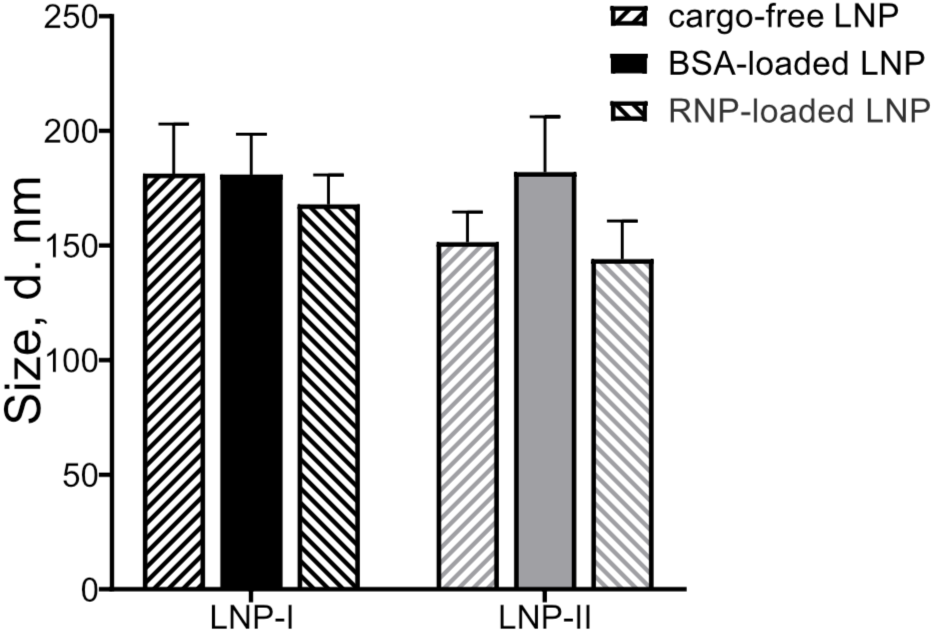
Comparison of the size of two LNP formulations encapsulating different cargos. Hydrodynamic diameters of LNP-I and LNP-II, either cargo-free, BSA-loaded, or RNP-loaded, were determined by DLS (intensity-weighted). Data are shown as mean ± SEM of 13 technical replicates for each formulation.

Encapsulation efficiency analysis showed that LNP-I and LNP-II had comparable efficiencies of 57.9% and 60.9%, respectively (Table 2). Efficiencies were calculated using Eq. 2, where C_UP_ represents the RNP concentration in the purified sample, and C_FULL_ represents the RNP concentration of the unpurified sample. Quantification was performed via reverse-phase HPLC using a previously established calibration curve and RNP standard chromatograms (Fig. S4A, B).

**Table 2.** Encapsulation efficacies of RNP in LNP-I and LNP-II.

| Composition | Encapsulation efficacy, % |
| --- | --- |
| LNP-I | 57.89 |
| LNP-II | 60.90 |

### LNPs deliver functional CasG and sgRNA ribonucleoprotein complexes to mammalian cells

Cas9-RNP delivery was assessed using LNP-I formulation, which demonstrated consistently high encapsulation and transfection efficacy. LNP-I was used to encapsulate functional Cas9 proteins and to test whether encapsulated proteins retained enzymatic activity. As LNPs are a usual delivery tool for various gene editing tools, we decided to test our proprietary LNPs by encapsulating widely used SpCas9 and sgRNA ribonucleoprotein (RNP) complex for a proof-of-principle assay (Fig. 6). First, we tested different Cas9-GFP RNP concentrations (9-35 nM) encapsulated in LNPs on HEK293T/17 cells cultured after transfection for various times (3, 6, or 24 hours) (Fig. 6A, B). Wide-field microscopy images revealed Cas9-GFP signal in the cell cytoplasm and the nuclei (Fig. 6A). Quantitative analysis revealed that, independently of the conditions, about half of the cells had Cas9-GFP inside the cytoplasm and/or the nuclei (Fig. 6B). Efficient transfection of Cas9-GFP RNPs was observed within three hours after the addition of LNPs. Interestingly, 24 hours after transfection, a statistically significantly lower number of nuclei exhibited GFP fluorescence with 9 nM RNP concentration, indicating that dosage optimization can be used to determine the preferred duration of RNP presence within the cells. After 24 h transfection, efficacy did not differ between 15 nM and 35 nM conditions; thus, the 15 nM concentration was used for further cell culture experiments, culturing cells for 24 hours after treatment to capture reliable gene editing results. To define the stability of RNP cargo encapsulated in LNP-I, prepared LNPs were either used fresh on the day of preparation or stored for 2 or 7 days at 4 °C before using them for cell treatment (Fig. 6C). We found that LNP-I were stable for at least a week, as there were no statistically significant differences in cell transfection efficacy for all the preparations used. Since LNP delivery of RNPs resulted in Cas9-GFP localization to the cell nuclei, we assessed the enzymatic activity of TrueCut v2 Cas9 encapsulated in LNP-I by performing the T7 endonuclease I assay (Fig. 6D, E). This universal assay detects indels in cells’ genomic DNA after cleavage by the Cas9 RNP. We targeted three benchmarking genes: *CDK4*, *RUNX1*, and *WTAP*, which are widely used in various CRISPR screens (30–32). As a reference, we used our proprietary LNPs alongside the commercial Lipofectamine CRISPRMAX transfection reagent, optimized for Cas9 RNP delivery. Heteroduplexes, formed due to genomic DNA indels, were detected after both LNP-I and CRISPRMAX delivery of Cas9 RNPs (Fig. 6D). Cleaved and non-cleaved DNA band intensities were plotted, and T7E1 cleavage efficacy was quantified, revealing a comparable T7E1 cleavage efficacy between our proprietary LNP-I and commercial lipofection reagent (Fig. 6E). Importantly, LNP-I delivery lead to a higher level of heteroduplexes at the *CDK4* site, confirming that LNP-I formulation can be used to encapsulate and deliver functionally active Cas9 RNPs at the same or better efficacy as commercially available alternatives.

**Fig. 6.**
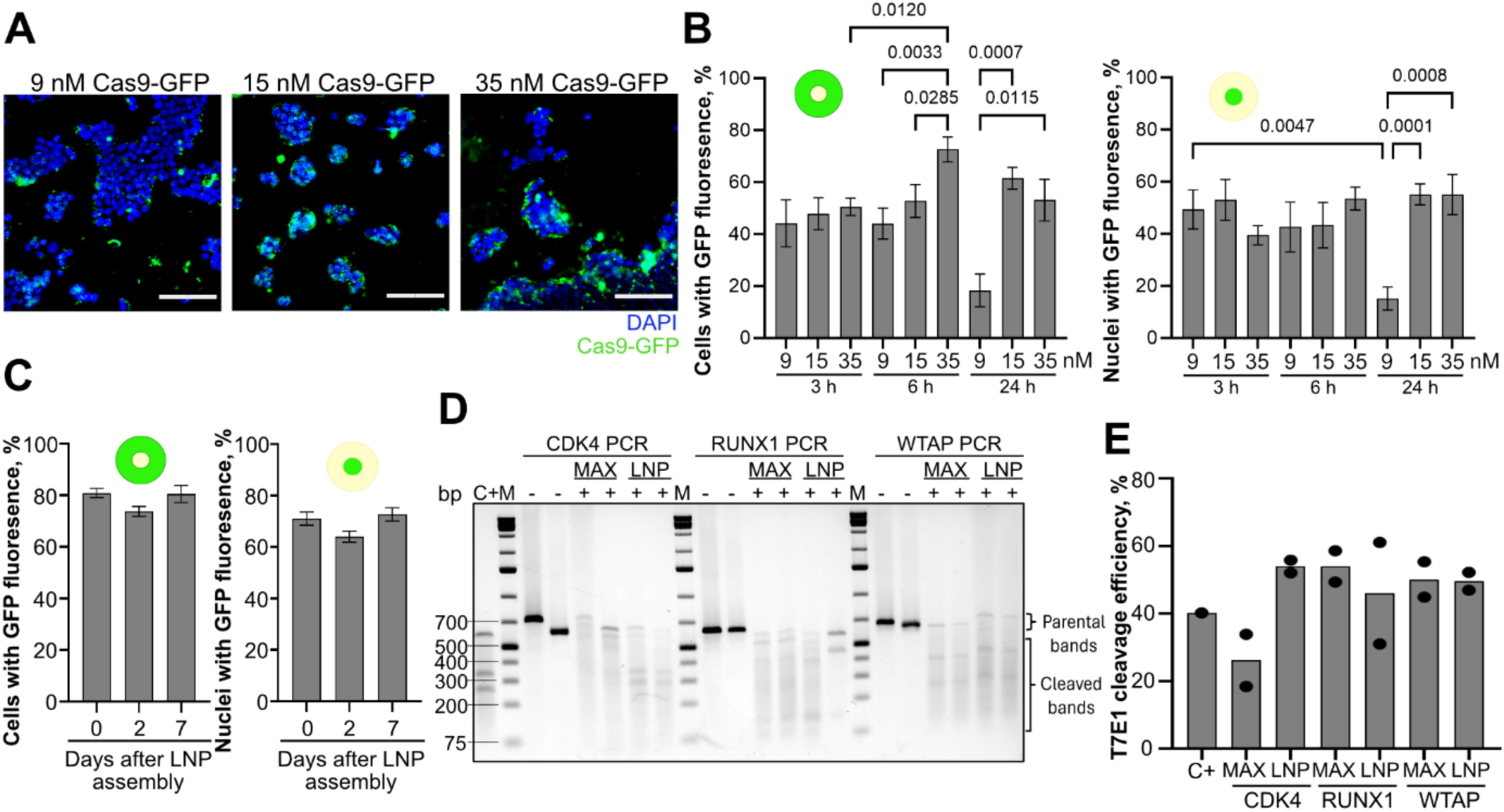
Functional delivery of CasG and sgRNA ribonucleoprotein complex to mammalian cells using LNP-I. **(A)** Widefield microscopy images of HEK293T/17 cells treated with Cas9-GFP RNP complexes encapsulated in LNP-I at varying concentrations (9 nM, 15 nM, 35 nM). Scale bar 100 µm. **(B)** Quantification of HEK293T cells showing GFP fluorescence in the cytoplasm or nucleus at different time points and LNP concentrations. **(C)** Quantification of HEK293T cells with cytoplasmic or nuclear GFP signal following treatment with LNPs of varying post-preparation time. **(D)** Functional assessment of TrueCut v2 Cas9 RNPs delivered via LNP-I (LNP) or commercial reagent Lipofectamine CRISPRMAX (MAX), using sgRNAs targeting *CDK4*, *RUNX1*, and *WTAP*, as assessed by T7 endonuclease I (T7E1) assay; **+/-** indicates whether RNPs were added to the cells. **(E)** Quantification of T7 endonuclease I cleavage efficacy to compare genome editing performance of Cas9 RNPs delivered by different methods. MAX – CRISPRMAX, LNP – LNP-I, C+ – positive T7E1 control. Data are presented as mean ± standard error of the mean (SEM) from 3 independent biological replicates (B, C), or as means with individual values from two technical replicates shown as dots (E). Means were compared by two-way ANOVA and *post-hoc* Holm-Šídák test (B) or one-way ANOVA and *post-hoc* Tukey‘s test (C). *p*-values < 0.05 were considered significant.

## Discussion

In this study, we developed proprietary LNPs with different lipid compositions (Fig.1). LNP-I showed the highest encapsulation efficacy of 68.2% (Table 1). Organotypic brain culture experiments revealed a strong influence of lipid composition on mammalian cell transfection characteristics (Fig. 2). These results are consistent with recent studies reporting that lipid components and their relative proportions are significant determinants of transfection efficacy and even define organ specificity (14, 33,34). We demonstrated that LNP-I and LNP-II are effective lipofection formulations for mammalian cell transfection, both in 2D and 3D tissue cultures (Fig. 2, 4, 6).

Stepwise processing of LNP-I and LNP-II highlighted the impact of cargo loading and purification on particle characteristics. While loading increased the initial LNPs’ size, subsequent extraction and ultracentrifugation steps effectively restored uniform size distributions of LNPs (Fig. 3A). They ensured long-term stability, as indicated by consistent PDI values during storage (Fig. 3B, C). LNPs are commonly used for various *in vivo* studies, usually to target the periphery after systemic delivery (14,35,36).

There is a lack of studies investigating the capacity of LNPs to permeate the blood-brain barrier *in vitro* before performing *in vivo* studies; therefore, we tested our proprietary LNP-I and LNP-II on an *in vitro* brain endothelial cell transcytosis model (Fig. 4A, B). HPLC fraction analysis and wide-field microscopy of the cells determined that LNP-I and LNP-II did not permeate the *in vitro* transcytosis barrier (Fig. 4C–G). Our results suggest that our proprietary LNPs have limited ability to reach the brain after systemic delivery; thus, confirming that the LNP-I and LNP-II formulations are safe for peripheral applications and do not pose potential side effects on the central nervous system. Our data demonstrate the proof-of-principle that an *in vitro* transcytosis model can be used to evaluate LNP formulations before undertaking animal studies, which are time-consuming and require ethical considerations.

The encapsulation of RNP complexes into LNPs did not significantly impact nanoparticle size, demonstrating that both LNP-I and LNP-II can accommodate large RNP without compromising their structural integrity (Fig. 5). Importantly, we demonstrated that LNP-I formulation can be used to encapsulate and deliver functional protein–RNA complexes in mammalian cells, such as SpCas9 and sgRNA RNPs, demonstrating their applicability as the delivery tool for rapidly developing gene editors (Fig. 6). With optimized encapsulated RNP delivery concentration, LNPs were stable for up to a week, further demonstrating their applicability for the delivery to mammalian cells (Fig. 6 A–C). Other studies suggest even longer shelf-life stability periods (37,38), but long-term studies are yet to be performed for the LNP-I formulation. Furthermore, with LNP-I, we successfully delivered functional RNPs that created indels in three targeted benchmarking genes, *WTAP*, *RUNX1*, and *CDK4* (Fig. 6D, E), in line with earlier findings demonstrating LNPs as an efficient delivery tool for highly functional RNPs (39–41).

In conclusion, this study demonstrates the potential of our proprietary LNP formulation to advance the characterization and application of lipid nanoparticles. These LNPs can deliver proteins or gene-editing RNPs in both 2D mammalian cell cultures and 3D tissue models, highlighting their versatility as a robust delivery system across diverse experimental platforms.

## Ethics approval

This study was conducted in accordance with Directive 2010/63/EU requirements and was approved by the Lithuanian State Food and Veterinary Service (Permit No G2-92, B6-(1.9)-2653).

## Availability of data and materials

All data generated or analysed during this study are included in this published article and its supplementary information files. Data sharing is not applicable to this article as no datasets were generated during the current study.

## Authors’ contributions

E.D.V., M.M., B.P., U.N. and R.B. designed the research, analyzed the data and wrote the paper. E.D.V.: investigation, validation, conceptualization, writing; M.M.: investigation, validation, conceptualization, writing; B.P.: methodology, data curation; N.D.: methodology, conceptualization; S.K.: methodology, conceptualization; C.Y.K.: conceptualization; Y.T.C.: conceptualization; U.N.: supervision, conceptualization, writing and editing, project administration; R.B.: supervision, conceptualization, writing and editing. All authors have read and agreed to the published version of the manuscript.

## Acknowledgements

The authors thank Evelina Jankaityte for the initial results on the preparation and formulation of lipid nanoparticles. We gratefully acknowledge the Sector of Amyloid Research for allowing us to use their microplate reader and the Department of Protein–DNA Interactions for providing access to the DLS instrument at the Vilnius University Life Sciences Center, Institute of Biotechnology.

