## Supplemental for "Improved Protein Encapsulation and Delivery by Lipid Nanoparticles with Refined Ionizable Lipid Content"

*Equal contributions

### Experimental Section

Notably, 1% (v/v) Triton X-100, as determined in prior experiments considering complete lipid nanoparticle destruction (Figure S1A) and acetonitrile/water elution buffer disrupts the lipid nanoparticles (Figure S2B).

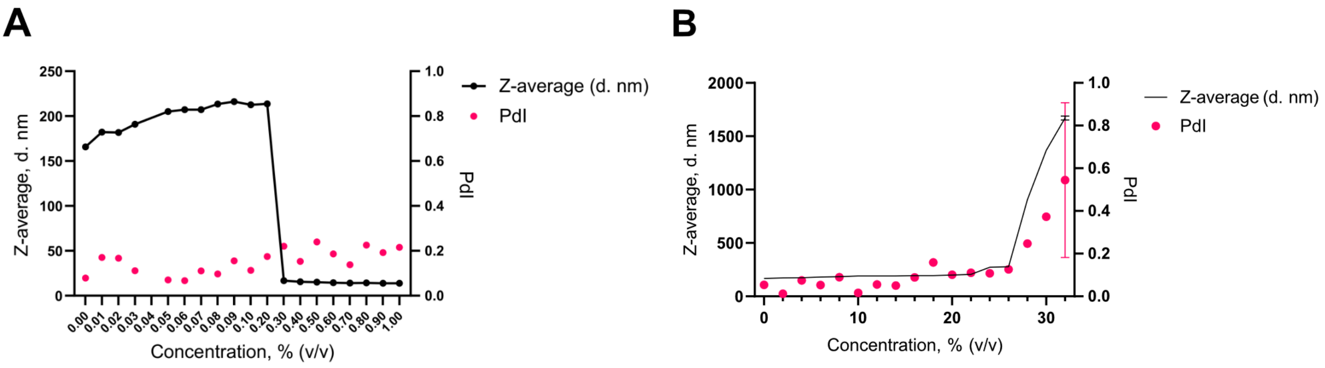


**Figure S1. (A)** Dependency of nanoparticle disruption on Triton X-100 concentration. Z-average diameters and polydispersity index (PDI) were monitored as a function of detergent concentration (v/v). Increasing Triton X-100 concentrations progressively destabilized the LNPs, with complete disruption achieved above ~0.5% (v/v). This established the minimal detergent concentration required for releasing encapsulated protein. **(B)** Dependency of nanoparticle disruption on acetonitrile concentration. Z-average diameters and polydispersity index (PDI) were monitored as a function of acetonitrile/water fraction (v/v).

BSA was chosen as a model of protein cargo due to its stability, accessibility, relatively low cost, and the ease of fluorescent detection after conjugation with AlexaFluor488. The calibration curve for BSA quantification by HPLC and representative chromatograms of BSA are shown in Figure S2 A,B.

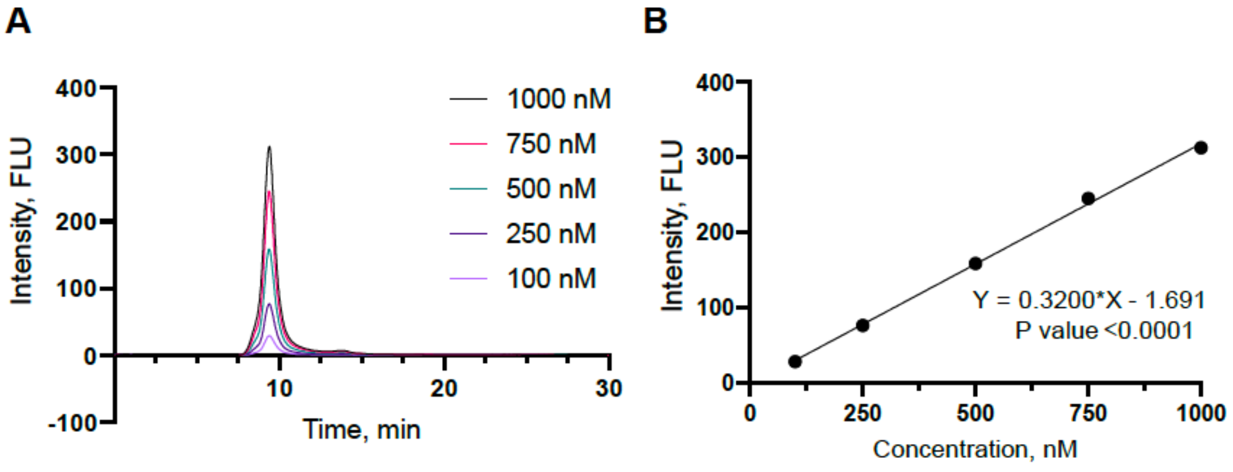


**Figure S2. (A)** Calibration curve for BSA quantification by HPLC. (**B)** Representative chromatograms of BSA standards (100–1000 nM) with the linear regression ($R^2 > 0.99$, $p < 0.0001$).

Representative chromatograms of BSA-loaded LNPs (I–IV) under different preparation conditions: intact nanoparticles (“full”), lysed nanoparticles (“full + Triton”), supernatant fractions (“up”), and detergent-treated supernatants (“up + Triton”). Distinct protein peaks were detected only in Triton-treated samples, confirming that detergent lysis is required to release encapsulated BSA for accurate quantification (Figure S3 A-D).

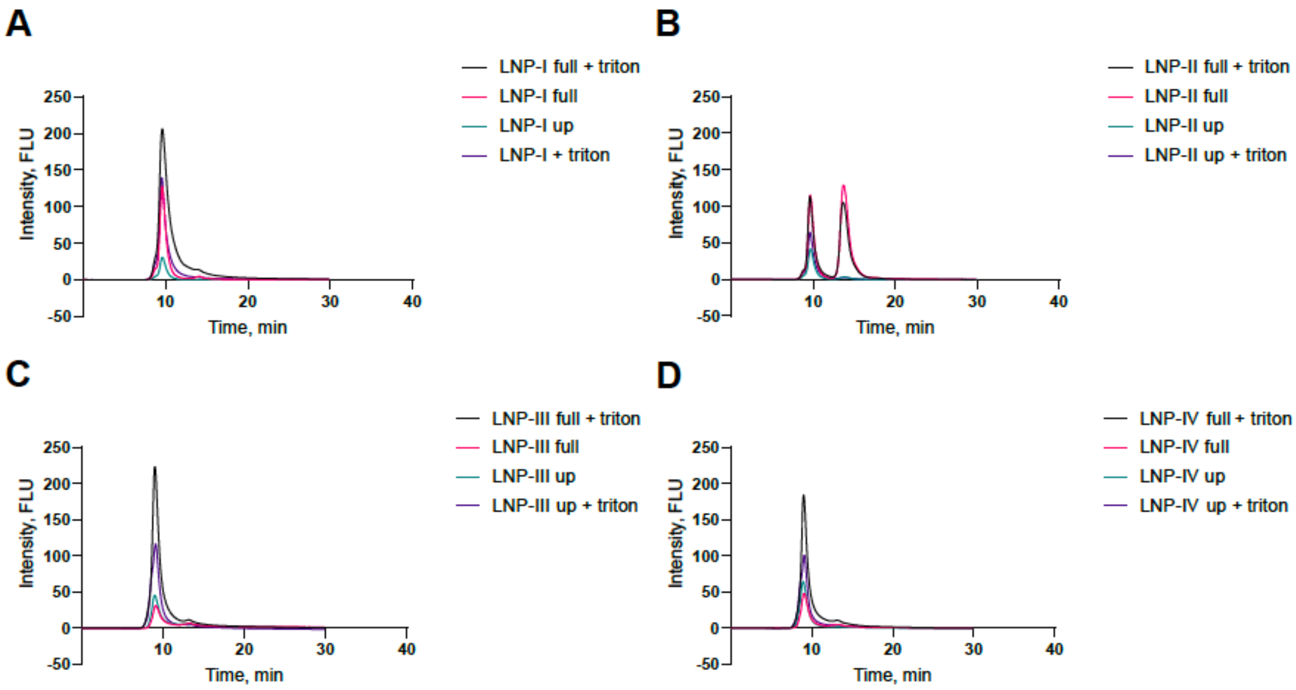


**Figure S3.** (A–D) Representative chromatograms of BSA-loaded LNPs (I–IV) under different preparation conditions: intact nanoparticles (“full”), lysed nanoparticles (“full + Triton”), supernatant fractions (“up”), and detergent-treated supernatants (“up + Triton”).

The **c**alibration curve for Cas9–sgRNA ribonucleoprotein (RNP) quantification by reverse phase HPLC and chromatograms of RNP standards (35–350 nM) resulting in a linear regression ($p = 0.0420$) are shown in Figure S4 A-B. For these measurements, detergent lysis was not necessary because the acetonitrile/water elution buffer used with the Brownlee Bio C18 column directly disrupted LNPs.

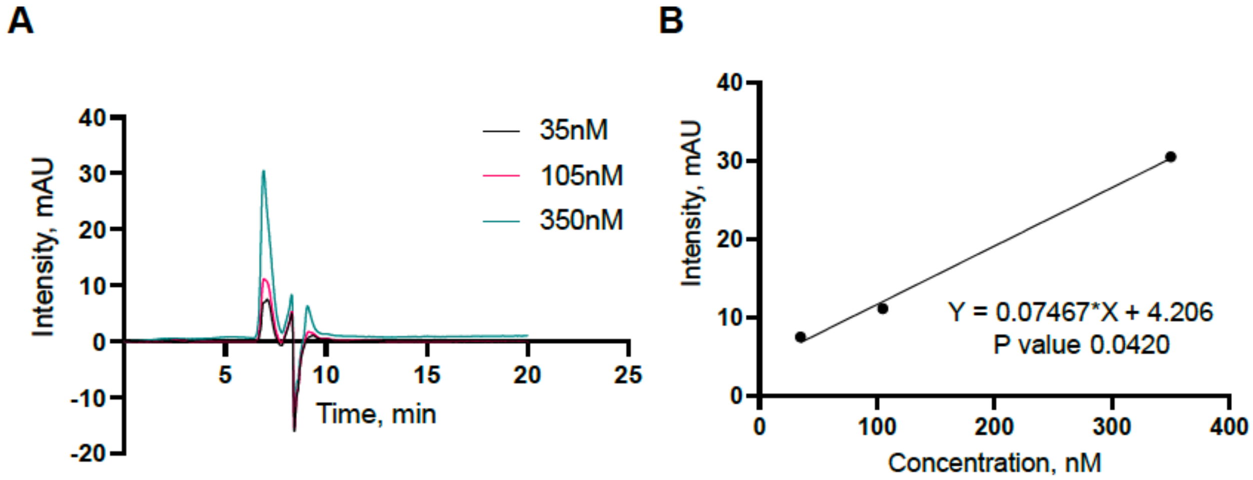


**Figure S4 (A)** Calibration curve for RNP quantification by reverse-phase HPLC. **(B)** Chromatograms of RNP standards (35–350 nM) with resulting linear regression ($p = 0.0420$).
